# Environmental modulation of social isolation-induced sleep loss in *Drosophila*

**DOI:** 10.64898/2026.09.23.753895

**Authors:** Brissa Castillo, Fiona Gugala, Yangyuan Li, Feda Hammood, Wanhe Li

## Abstract

Chronic social isolation induces persistent behavioral and physiological changes across species. In humans, prolonged social isolation is associated with insomnia and increased risks of cognitive decline, cardiovascular disease, compromised immunity, and premature mortality. Yet whether these consequences are modified by the environment in which social isolation is experienced remains unclear. In particular, can access to social sensory information from conspecifics alleviate the effects of isolation and can various ambient environmental settings modify them? A related question arises during behavioral measurement because behavioral assays themselves impose a new environmental context. Whether these assays simply capture the enduring effects of prior social experience or further modify those effects remains unknown. Using social isolation-induced sleep loss in *Drosophila* as a model, we systematically examined how the social, spatial, and photoperiodic environments during isolation and behavioral monitoring influence the isolation phenotype. Providing isolated flies with visual, olfactory, or tactile social cues from conspecifics living behind a divider, either individually or in combination, was insufficient to alleviate sleep loss. Spatial confinement exacerbated isolation-induced sleep loss, while winter-like photoperiod in spatially confined *Drosophila* Activity Monitor assays revealed the clearest progressive decline in sleep during sleep monitoring. Together, these findings demonstrate that social isolation-induced sleep loss is both enduring and dynamic: it persists despite exposure to social cues yet remains sensitive to the environmental context in which isolation is experienced and behavior is measured. More broadly, our study establishes a tractable framework for investigating the environmental modulation of chronic social isolation.

## Introduction

Social isolation produces profound behavioral and physiological effects across species and is increasingly recognized as an emerging public health concern [1–8]. These effects are primarily attributed to the lack of social interaction. However, social isolation does not occur in an environmental vacuum: isolated individuals may receive social sensory information and interact with ambient conditions that may change over time. Whether these environmental features alleviate, exacerbate, or alter the progression of isolation-induced behavioral changes remains largely unexplored.

Behavioral assays provide the foundation for understanding how experience shapes brain function [9–11]. To enable precise and reproducible measurements, behavioral phenotypes are typically characterized by transferring animals under defined treatment conditions into simplified experimental settings, such as individual chambers, confined arenas, or standardized testing apparatuses. These treatment and assay conditions are generally assumed to provide a neutral process through which the behavioral consequences of prior experiences can be measured as endpoints. However, because treatment conditions can vary and behavioral measurements inevitably alter the environment experienced by the animal, important questions considering these environments remain largely unexplored: can the treatment environment modify behavior, and can the measurement environment itself continue to influence the behavioral process under investigation?

Measuring sleep changes induced by social isolation presents a compelling example of these challenges. We and others have shown that chronic social isolation robustly reduces sleep in *Drosophila* [8, 12–17]. In these experiments, sleep is typically recorded from individually housed animals following prolonged group housing or social isolation in standard food vials, demonstrating that prior social experience has persistent effects on sleep. Similarly, sleep and many other behaviors in rodents and humans are often assessed after subjects have experienced defined treatment conditions and are subsequently transferred to standardized settings for individual testing. Because animals with different social histories are ultimately placed into the same isolated recording environment, it remains unclear whether behavioral monitoring simply captures the endpoint of prior social experience or allows the effects of social isolation to continue developing during measurement.

Here, we address these questions by systematically investigating how the social, spatial, and photoperiodic environments experienced during isolation and behavioral monitoring influence social isolation-induced sleep loss in *Drosophila*. Our findings reveal that isolation-induced sleep loss is both enduring and dynamic: it persists despite access to social sensory cues but remains strongly influenced by the environmental context in which isolation is experienced and behavior is measured.

## Results

### Social cues are insufficient to alleviate chronic social isolation-induced sleep loss

To determine whether social environment could alleviate the behavioral consequences of chronic social isolation, we exposed isolated *Drosophila melanogaster* to defined sensory cues from conspecifics during a 7-day treatment period following a 3-5days socialization phase under standard group housing conditions [8] **(Fig. 1A).** We previously found that, after 7 days of social isolation, flies exhibit reduced total and daytime sleep, including reduced sleep during the first four hours of the light phase (ZT0-4) of a standard 12:12 LD (12hr light and 12hr dark) cycle, with an increased number of sleep bouts and a shorter average bout length, indicating fragmented sleep. We refer to this phenotype as chronic social isolation-induced sleep-loss thereafter in this work. Although we and others previously observed isolation-induced sleep loss in both male and female flies, we focused the present study on males to enable systematic comparison across the large number of social, sensory, and environmental conditions examined. We systematically tested whether olfactory, visual, and mechanosensory cues, either individually or in combination, could rescue the sleep-loss phenotype associated with chronic isolation **(Fig. 1B and Fig. S1).**

**Figure 1.**
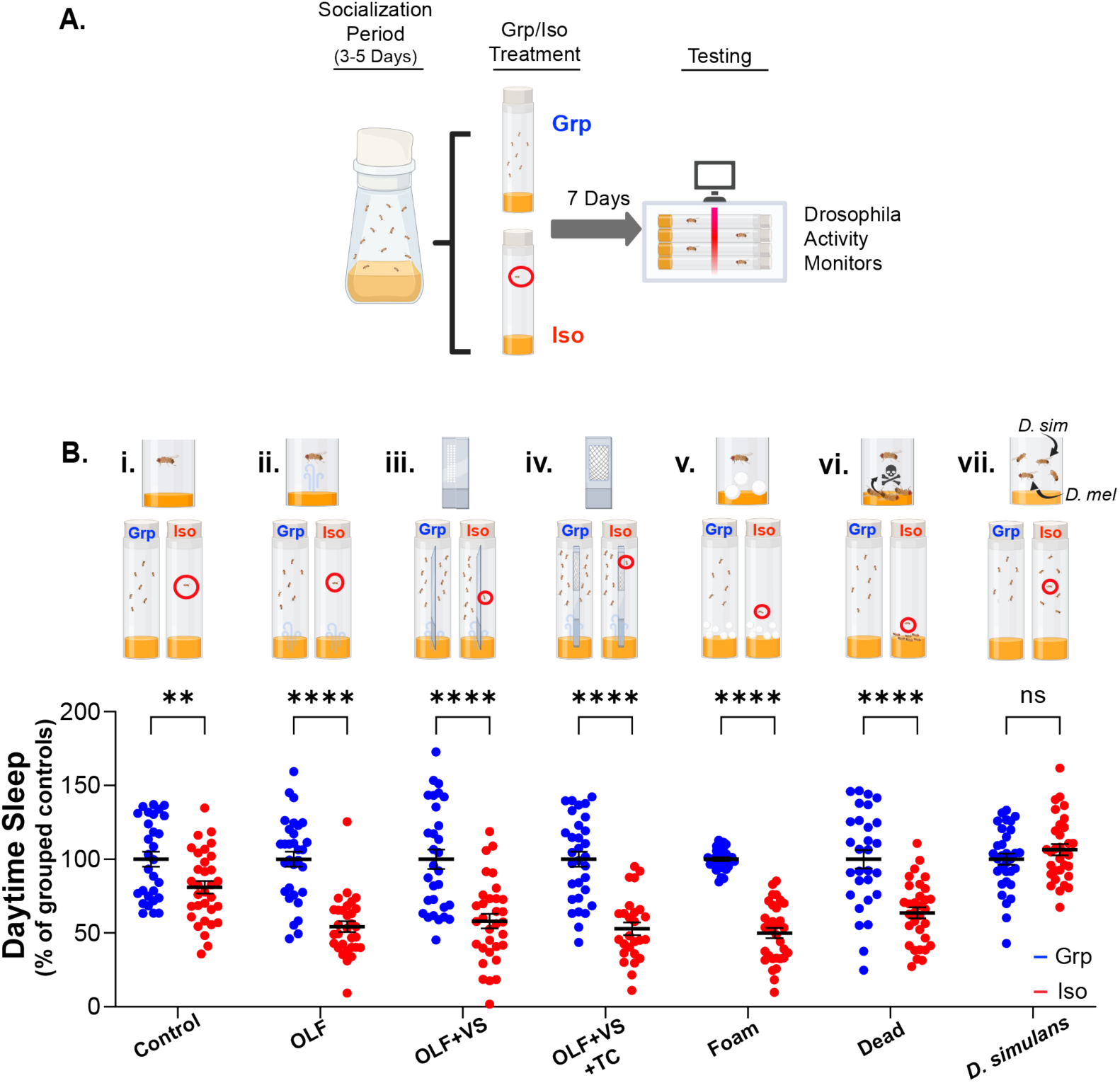
Social cues are insufficient to alleviate chronic social isolation-induced sleep loss. (A) After eclosion, flies were collected and kept in groups for 3–5 days before being assigned to either 7 days of group housing or 7 days of chronic social isolation. (B) Schematic of experimental conditions used to provide isolated and grouped wild-type flies (i) and with sensory cues from conspecifics: (ii) vials pre-conditioned by other flies, leaving olfactory cues prior to introduction of experimental flies; (iii) a perforated divider allowing olfactory and visual cues to be communicated to experimental flies; (iv) a divider with a mesh-screened window allowing experimental flies to receive olfactory, visual, and tactile cues from conspecifics (See Supplemental Video 1); (v) exposure to foam balls; (vi) exposure to dead flies; and (vii) exposure to a heterospecific species, D *simulans*. Quantification of daytime sleep expressed as a percentage of grouped controls, normalized to 100% (mean ± s.e.m.; individual data points shown). See Figure S1 for total sleep, daytime sleep, nighttime sleep, ZT0-4 sleep, as well as daytime and nighttime bout number and bout duration from the same dataset. Statistical analysis was performed using two-sided unpaired t-tests with Welch’s correction. \**P* < 0.05; \*\**P* < 0.01; \*\*\**P* < 0.001; \*\*\*\**P* < 0.0001; NS, not significant; *n* = 29–32 flies per condition.

We first examined whether olfactory cues alone were sufficient to mitigate isolation-induced sleep loss using odor-conditioned vials **(Fig. 1B-ii)**. Wild-type flies were used to condition the vials. They were housed in groups for 3 days to deposit residual chemical and pheromonal cues before experimental flies were introduced. Despite exposure to these socially conditioned environments, isolated flies continued to exhibit reduced sleep comparable to fully isolated controls, whereas grouped flies maintained normal sleep levels. Thus, residual olfactory cues from conspecifics are insufficient to rescue social isolation-induced sleep loss.

We next assessed whether combined olfactory and visual cues could alleviate the isolation-induced sleep-loss phenotype using a transparent perforated divider that permitted the isolated fly to see and smell other flies on the other side of the divider while preventing direct physical interaction **(Fig. 1B-iii)**. Under these conditions, isolated flies continued to display reduced sleep, demonstrating that access to visual and olfactory social information does not compensate for the absence of direct interaction.

To determine whether adding mechanosensory input was sufficient to alleviate isolation-induced sleep loss, we introduced a mesh-divider paradigm that allowed some tactile contact in addition to visual and olfactory communication between the isolated fly and other flies on the other side of the divider **(Fig. 1B-iv, Fig. 1C and Supplemental Video 1)**. Despite access to all three sensory modalities, isolated flies maintained reduced sleep levels similar to that of fully isolated animals, whereas grouped flies remained unaffected. Together, these findings demonstrate that multimodal sensory exposure alone is insufficient to reverse chronic isolation-induced sleep loss.

Next, we tested whether more complex social stimuli could influence the isolation phenotype by exposing isolated *D.melanogaster* flies to small foam balls **(Fig. 1B-v)**, dead *D.melanogaster* conspecifics **(Fig. 1B-vi)**, or live *D. simulans* partners **(Fig. 1B-vii)**. Neither foam balls nor dead conspecifics restored sleep in isolated flies. In contrast, housing isolated *D. melanogaster* males with live *D. simulans* partners restored sleep to levels comparable to socially grouped controls. These findings demonstrate that the sensory manipulations tested cannot substitute for interaction with a living social partner in alleviating chronic social isolation-induced sleep loss.

### Spatial restriction exacerbates chronic social isolation-induced sleep loss

We next asked whether the spatial environment could modify the isolation-induced phenotype. We housed male *D.melanogaster* under two distinct spatial conditions: a standard fly food vial environment or a custom-designed housing system (a box with 96 individual fly food wells) with an ∼85% reduction in available space **(Fig. 2A)**. Within each housing condition, flies were maintained either in groups or under isolation for a duration of 7 days before measuring their sleep in the *Drosophila* Activity Monitor (DAM) assay. This experiment revealed that social isolation-induced sleep-loss was further enhanced when isolated flies were maintained under spatially restricted conditions.

**Figure 2.**
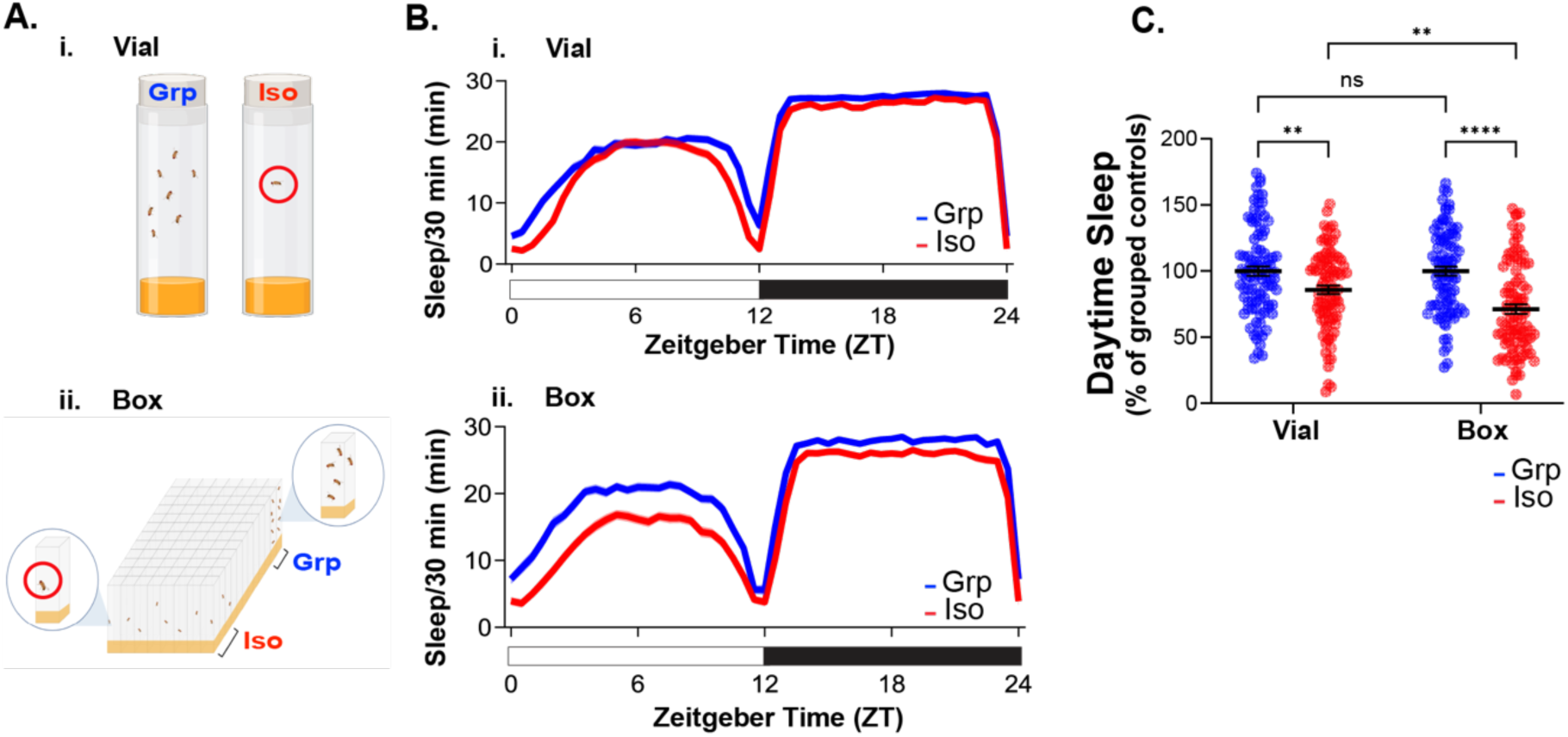
Spatial restriction exacerbates chronic social isolation-induced sleep loss. (A) Schematics of experimental conditions used to house grouped and isolated flies in environments with different spatial constraints: (i) traditional vial-based housing; (ii) a custom-designed box-based housing system providing approximately 85% reduction in living space. (B) Sleep profiles of grouped and chronically isolated flies under vial-based and box-based housing conditions. (C) Quantification of daytime sleep expressed as a percentage of grouped controls, normalized to 100% (mean ± s.e.m.; individual data points shown). See Figure S2 for quantification of all sleep parameters from the same dataset. Statistical analysis was performed using two-way ANOVA followed by Fisher’s LSD multiple-comparison tests. A significant interaction was detected between housing conditions and social state (*P* = 0.0337). Isolated flies exhibited reduced sleep compared with grouped flies under both vial-based housing and spatially restricted housing. Housing conditions did not significantly affect sleep in grouped flies but significantly affected sleep in isolated flies. See Figure S2 for quantification of all sleep parameters from the same dataset. \**P* < 0.05; \*\**P* < 0.01; \*\*\**P* < 0.001; \*\*\*\**P* < 0.0001; NS, not significant; n = 86–92 flies per condition.

Importantly, spatial restriction alone did not alter sleep in grouped flies, as socially housed animals displayed comparable sleep levels between vial and reduced-space environments **(Fig. 2C and Fig. S2)**. These findings demonstrate that the physical environment in which isolation is experienced can modulate the severity of the resulting behavioral phenotype.

### Diverse photoperiods and light histories do not eliminate chronic social isolation-induced sleep loss

We next examined the light environment. Because environmental light strongly influences sleep regulation and circadian timing, we asked whether photoperiod or prior light exposure could modify the isolation-induced phenotype. Following a standardized socialization period, flies were maintained under grouped or isolated conditions and exposed to distinct photoperiod environments, including equinox-like 12:12 LD, winter-like 08:16 LD (8hr light and 16hr dark), summer-like 16:08 LD (16hr light and 8hr dark) **(Fig. 3A, Fig. S3 and Fig. S4)**, constant light (LL), and constant dark (DD) conditions **(Fig. S5)**. Sleep was then recorded using DAM under the corresponding treatment conditions. Across all tested photoperiods, chronically isolated animals consistently exhibited reduced daytime sleep compared with their grouped controls **(Fig. S5Eiii-iv)**.

**Figure 3.**
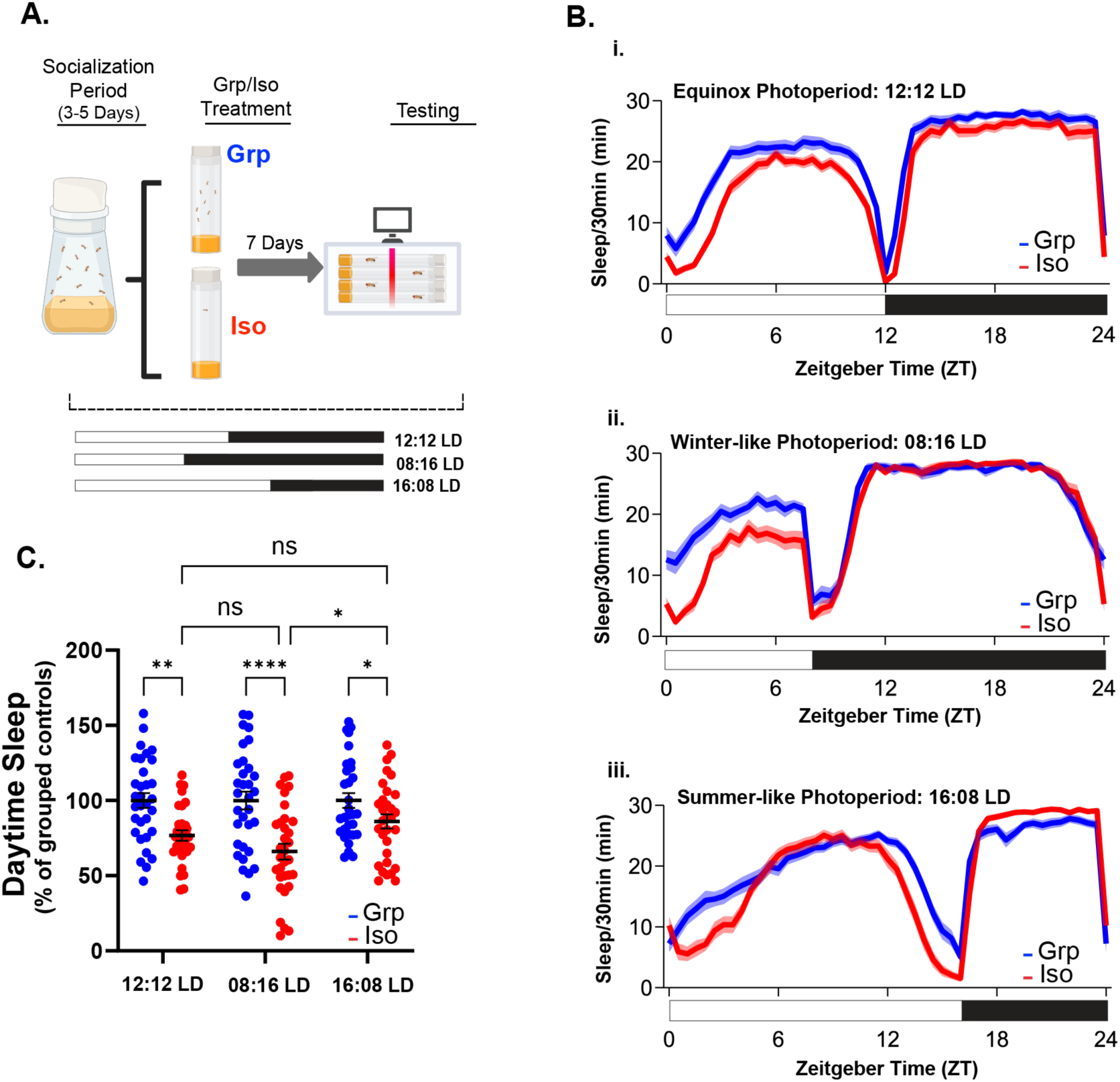
Winter-like photoperiod enhances chronic social isolation-induced sleep loss. (A) Schematic of experimental design. Following a socialization period, flies were assigned to grouped or isolated conditions and maintained under equinox-like (12:12 LD), winter-like (08:16 LD), and summer-like (16:08 LD) photoperiods at 21.5°C. (B) Sleep profiles of grouped and chronically isolated flies under each condition: (i) 12:12 LD, (ii) 08:16 LD, and (iii) 16:08 LD. (C) Quantification of daytime sleep expressed as a percentage of the corresponding grouped controls, normalized to 100% (mean ± s.e.m.; individual data points shown). See Figure S3 for quantification of all sleep parameters from the same dataset. Statistical analysis was performed using two-way ANOVA followed by Tukey’s multiple-comparison tests. Isolated flies exhibited reduced daytime sleep compared with grouped flies under all three photoperiods, with a greater reduction under 08:16 LD than under 16:08 LD (*P* = 0.0125). Significance thresholds: \**P* < 0.05; \*\**P* < 0.01; \*\*\**P* < 0.001; \*\*\*\**P* < 0.0001; NS, not significant. n = 31–32 flies per condition.

The persistence of chronic social isolation-induced sleep loss under LL and DD conditions further indicates that the phenotype does not require normal circadian rhythmicity or visual perception of social isolation to be expressed, as constant light disrupts circadian rhythmicity through light-dependent TIMELESS degradation, whereas constant darkness does not allow the isolated flies to visually perceive that they are alone. To distinguish the effects of current light conditions from prior environmental light conditions, flies were also recorded using DAM following transitions of the light conditions, including LD→LD, LL→LD, and DD→LD **(Fig. S6A-C)**. Regardless of the previous light environment or transition paradigm, isolated flies continued to exhibit reduced sleep compared with grouped controls **(Fig. S6AD)**.

These results demonstrate that social isolation-induced sleep loss persists across diverse environmental light conditions and light histories. Interestingly, the magnitude of social isolation-induced sleep loss appeared to be modulated by photoperiod. During the light phase, winter-like 08:16 LD conditions produced the strongest phenotype, with isolated flies exhibiting approximately 50% less daytime sleep than grouped controls, whereas 12:12 LD produced an intermediate effect and summer-like 16:08 LD the weakest effect **(Fig.3C)**.

### Spatial restriction and winter-like photoperiod reveal progressive social isolation-induced sleep loss

Our environmental manipulations identified spatial restriction as a condition that exacerbated chronic social isolation-induced sleep loss, while winter-like photoperiod produced the largest reduction in sleep among the photoperiods tested. This observation led us to reconsider the conditions under which sleep is conventionally measured. In the standard DAM assay, individual flies are confined within narrow tubes, creating an isolation experience in a spatially restricted environment. We therefore asked whether using the DAM assay under a winter-like photoperiod condition could reveal changes in sleep that develop progressively from the onset of individual behavioral monitoring.

In the next experiment, we eliminated the 7-day chronic social isolation period from our paradigm. Instead, socially experienced flies were transferred directly from group housing in large food bottles into individual, spatially restricted cuvettes and monitored using DAM. Therefore, the beginning of behavioral recording also marked the beginning of social isolation. During the recording period, flies were maintained under equinox-like 12:12 LD, winter-like 08:16 LD, or summer-like 16:08 LD conditions **(Fig. 4A)**.

**Figure 4.**
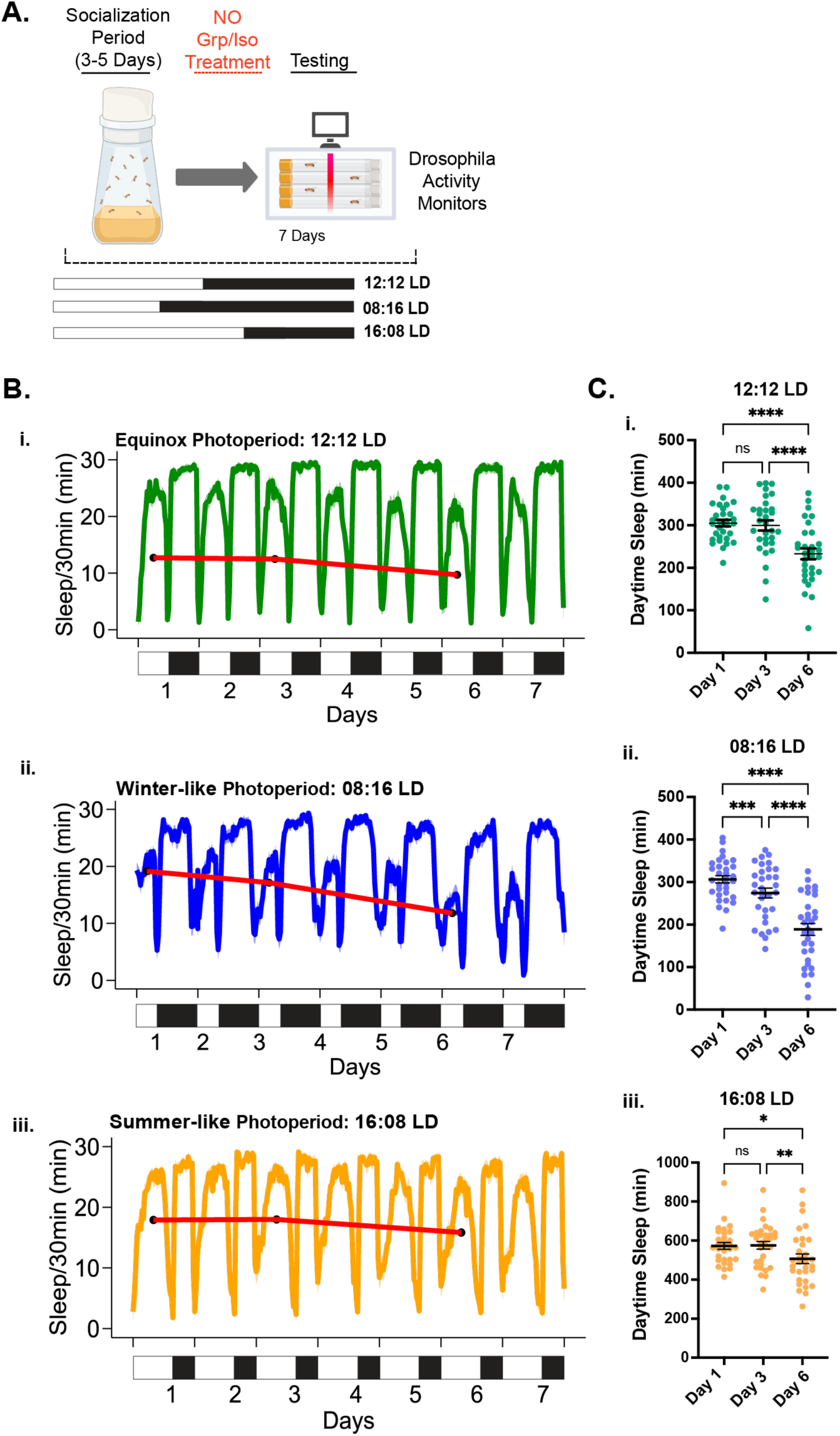
Spatial restriction and winter-like photoperiod reveal progressive social isolation-induced sleep loss. (A) Schematic of the experimental design. Following a socialization period, flies were transferred directly into individual glass tubes of the *Drosophila* Activity Monitor (DAM) and maintained under equinox-like (12:12 LD), winter-like (08:16 LD), or summer-like (16:08 LD) photoperiods at 21.5 °C. (B) Sleep profiles across successive days of isolation under (i) 12:12 LD, (ii) 08:16 LD, and (iii) 16:08 LD conditions. Red lines indicate changes in average daytime sleep (per 30min) on Days 1, 3, and 6. (C) Quantification of daytime sleep on Days 1, 3, and 6 under each photoperiod condition (mean ± s.e.m.; individual data points shown). Statistical analysis was performed using repeated-measures ANOVA with Geisser–Greenhouse correction followed by Tukey’s multiple-comparison tests. A significant effect of time was detected under all photoperiod conditions (*P* < 0.0001). Under 12:12 LD and 16:08 LD, daytime sleep did not differ significantly between Days 1 and 3 but decreased significantly by Day 6. Under 08:16 LD, daytime sleep was significantly reduced by Day 3 and declined further by Day 6. \**P* < 0.05; \*\**P* < 0.01; \*\*\**P* < 0.001; \*\*\*\**P* < 0.0001; NS, not significant; n = 30-32 flies per condition.

Continuous sleep profiling revealed a striking progressive reduction in daytime sleep under winter-like 08:16 LD conditions **(Fig. 4B-ii)**. In comparison, this progression was less pronounced under 12:12 LD and summer-like 16:08 LD conditions **(Fig. 4B-i, iii)**. Under these two photoperiods, daytime sleep remained relatively stable during the early recording period and declined only later around Day 6 In contrast, under winter-like 08:16 LD condition, daytime sleep was significantly reduced by Day 3 relative to Day1 and continued to decline through Day 6 **(Fig. 4C-ii)**. This experiment demonstrated that the spatially restricted DAM environment, particularly under winter-like photoperiod, can reveal the progressive development of social isolation-induced sleep loss during behavioral monitoring.

## Discussion

This study demonstrates that chronic social isolation in *Drosophila melanogaster* produces a persistent sleep-loss phenotype that is largely resistant to modulation by social sensory cues. Isolated flies exhibited robust sleep loss even when provided with olfactory, visual, and tactile information from conspecifics, either individually or in combination. This conclusion is further supported by our accompanying study showing that early-life social isolation primes animals for more aggressive neurodegenerative phenotypes: accelerated TDP-43 pathology and reduced lifespan. The same sensory manipulations also failed to rescue this phenotype. Importantly, the exacerbation of neurodegeneration was not explained by isolation-induced sleep loss, indicating that sleep loss and accelerated neurodegeneration are distinct consequences of chronic social isolation (see accompanying study by MurthyGowda et al) [18]. Together, these findings suggest that chronic social isolation establishes a persistent state with multiple downstream functional biological impacts that cannot be reversed by providing partial sensory features of social interaction.

Previous work showed that blind and anosmic mutant flies do not exhibit the normal increase in sleep following social enrichment [13]. Therefore, using sensory-deficient mutant animals in our paradigm would produce a potential ambiguity in interpretating the result. For example, an absence of sleep differences between grouped and isolated mutant animals could indicate that the disrupted sensory modality is required to perceive social isolation. Alternatively, it could reflect the mutants’ inability to interact normally with conspecifics and acquire the social experience required for normal level of sleep and other behavior in grouped controls [14, 19, 20]. In addition, it has been showed that social experience and pheromone-receptor activity jointly reshape transcriptional programs in olfactory sensory neurons and modulate courtship-related behavior [21, 22]. Our divider-based assays addressed this ambiguity by using wild-type animals while selectively controlling their access to visual, olfactory, and tactile cues from conspecifics. This design allowed us to conclude that these social cues, either individually or in combination, are insufficient to rescue isolation-induced sleep loss. In contrast, recent work in mice showed that touch-like tactile stimulation attenuates isolation-induced social rebound [23], suggesting that the ability of individual sensory cues to modify the effects of isolation may depend on the species and behavioral outcome examined.

Spatial restriction exacerbated chronic social isolation-induced sleep loss. This finding is particularly intriguing because the hDeltaK (P2) neurons in the central complex have previously been implicated in the response to chronic social isolation [8, 24]. Silencing these neurons during chronic social isolation abolished the associated sleep loss. hDeltaK neurons also signal the fly’s heading and locomotor speed. Recent studies further demonstrated a role for hDeltaK in goal-directed spatial navigation [25]. These neurons may therefore link the perception of spatial environment to the expression of isolation-induced sleep loss, although their role in mediating the effect of spatial restriction remains to be tested.

Collectively, our results suggest that chronic social isolation in *Drosophila* is encoded as a persistent internal state that cannot be readily reversed by sensory or environmental manipulations. However, this state is not immutable, as the conditions experienced during behavioral monitoring can shape the progression and expression of the isolation-induced phenotype. Under winter-like short photoperiod conditions, DAM monitoring revealed a progressive loss of sleep, demonstrating that behavioral assays can capture an evolving phenotype rather than just measure an endpoint. This interaction between social isolation and photoperiod may also have relevance to seasonal affective disorder, in which seasonal changes in light exposure are associated with altered sleep, circadian dysfunction, and mental health issues [26]. In *Drosophila*, recent studies have begun to define the molecular mechanisms by which the circadian system integrates photoperiodic and temperature cues to regulate seasonal adaptation [27–29]. Although the fly phenotype is not a direct model of this human disorder, it may provide a platform for studying how social isolation and seasonal conditions interact to influence sleep.

## Materials and Methods

### Fly culture

*Drosophila melanogaster* (wild-type Canton-S strain) were maintained on standard cornmeal–yeast–molasses–agar medium at 21.5 °C under a 12 h light:12 h dark (LD) cycle. Newly eclosed flies were collected and transferred to bottles, where males and females were allowed to socially interact for 3–5 days prior to experimental assignment.

### Chronic social isolation and group housing conditions

Following the socialization period, male flies were assigned to either chronic isolation or group-housing conditions using both a traditional vial-based paradigm (Li et al., 2021) and a newly developed box-based paradigm.

### Vial-based paradigm

Using the standard fly food vials (VWR 75813-166), flies were housed either individually (one fly per vial) or in groups of 25 flies per vial. Flies remained in their assigned condition for 7 days with approximately 56 cm³ of living space.

### Box-Based Paradigm

Individual flies were housed in separate compartments of a 96-well plate (VWR 76924-056), whereas grouped controls were housed at five flies per compartment for the same duration. This system reduced the available space per fly by approximately 85% compared with the traditional vial-based paradigm.

### Sensory cue manipulations using dividers

To examine the contributions of specific sensory modalities, we introduced dividers with different configurations into the vial-based housing system.

### Olfactory cues only

To generate an odor-conditioned environment, a mixed-sex group of flies was housed in a vial for 3 days to deposit olfactory cues. These flies were then removed, and experimental flies, either isolated individuals or groups of 25, were introduced for a 7-day treatment period.

### Olfactory and visual cues

To permit visual and olfactory interactions, a transparent plastic divider with small perforations was inserted into the vial, allowing flies housed on opposite sides to see one another and exchange olfactory cues while preventing physical contact. The perforations are too small for flies to pass through.

### Combined cues: olfactory, visual, and tactile cues

To permit visual, olfactory, and tactile interactions, a transparent plastic divider containing a mesh-screened window was inserted into the vial, allowing flies housed on opposite sides to see one another, exchange olfactory cues, and engage in some physical contact through the mesh. To verify that tactile interactions occurred across the mesh interface, the isolated and grouped flies on opposite sides of the divider were continuously recorded for 24 h using two cameras to capture instances of physical contact.

### Addition of foam objects

Small foam balls were added to both isolated and grouped housing conditions to assess whether the presence of non-social physical objects influenced behavioral outcomes associated with chronic isolation.

### Addition of dead flies

Wild-type flies were euthanized at −80 °C for 15 min, and 24 dead flies were introduced into each isolated and grouped vial to determine whether the presence of dead conspecifics alleviated isolation-induced sleep loss.

### Addition of Drosophila simulans

To assess whether heterospecific social interactions alleviated isolation-induced sleep loss, 24 *D. simulans* flies were introduced into each vial containing either isolated or grouped *D. melanogaster*. In this setup, *D. melanogaster* flies were either kept in isolation or grouped exclusively with *D. simulans*, with no conspecifics present.

### Locomotor Activity Assay

Locomotor activity was measured using the *Drosophila* Activity Monitoring (DAM) System (TriKinetics, Inc.). Individual flies were loaded into glass tubes with food, and activity was recorded in 1-min bins for consecutive light–dark (LD) cycles (or other lighting conditions as specified), beginning on the day after loading. Sleep parameters, including total sleep, daytime sleep, nighttime sleep, ZT0-4 sleep, sleep profiles, and sleep-bouts, were analyzed in R (v3.6) using custom code based on the rethomics package and visualized using GraphPad Prism.

### Quantification and Statistical Analysis

Statistical analyses were performed using GraphPad Prism. Experiments were independently repeated at least three times, and statistical comparisons were made between experimental groups tested in parallel within the same experiment. Representative datasets from one independent experiment are shown for each condition, unless otherwise indicated. Data are presented as mean ± s.e.m., with individual data points shown where indicated. Comparisons between two groups were performed using two-sided unpaired t-tests with Welch’s correction. Experiments involving two independent variables were analyzed using two-way ANOVA followed by Fisher’s LSD or Tukey’s multiple-comparison tests, as indicated in the corresponding figure legends. Repeated measurements across successive days were analyzed using repeated-measures ANOVA with Geisser–Greenhouse correction followed by Tukey’s multiple-comparison tests. Statistical significance was defined as \**P* < 0.05; \*\**P* < 0.01; \*\*\**P* < 0.001; \*\*\*\**P* < 0.0001; NS, not significant. Sample sizes (n) indicate the number of individual flies and are reported in the corresponding figure legends.

## Acknowledgments

This work was supported by the National Institute of General Medical Sciences (GM150832 to W.L.). We thank Min Feng, Swetha MurhyGowda, and Josh Dubnau for comments on the manuscripts. We thank Esther Doria and Min Feng for the isolation box design. We thank members of the Li laboratory for technical support. B.C. was supported by a graduate research fellowship from the Hagler Institute for Advanced Study at Texas A&M University. F.H. was supported by the Yalow Scholars Program and the McNulty Scholars Program at Hunter College. W.L. is a CPRIT Scholar in Cancer Research (Cancer Prevention and Research Institute of Texas, RR220021).

## Author Contributions

B.C. and W.L. conceived and designed research; B.C., F.G., and F.H. performed the experiments and conducted statistical analyses; Y.L. produced the box and divider tools; B.C. and W.L. wrote the manuscript.

## Supporting Information

**Supplemental Figures S1-S6**

**Supplemental Video 1**

**Figure S1.**
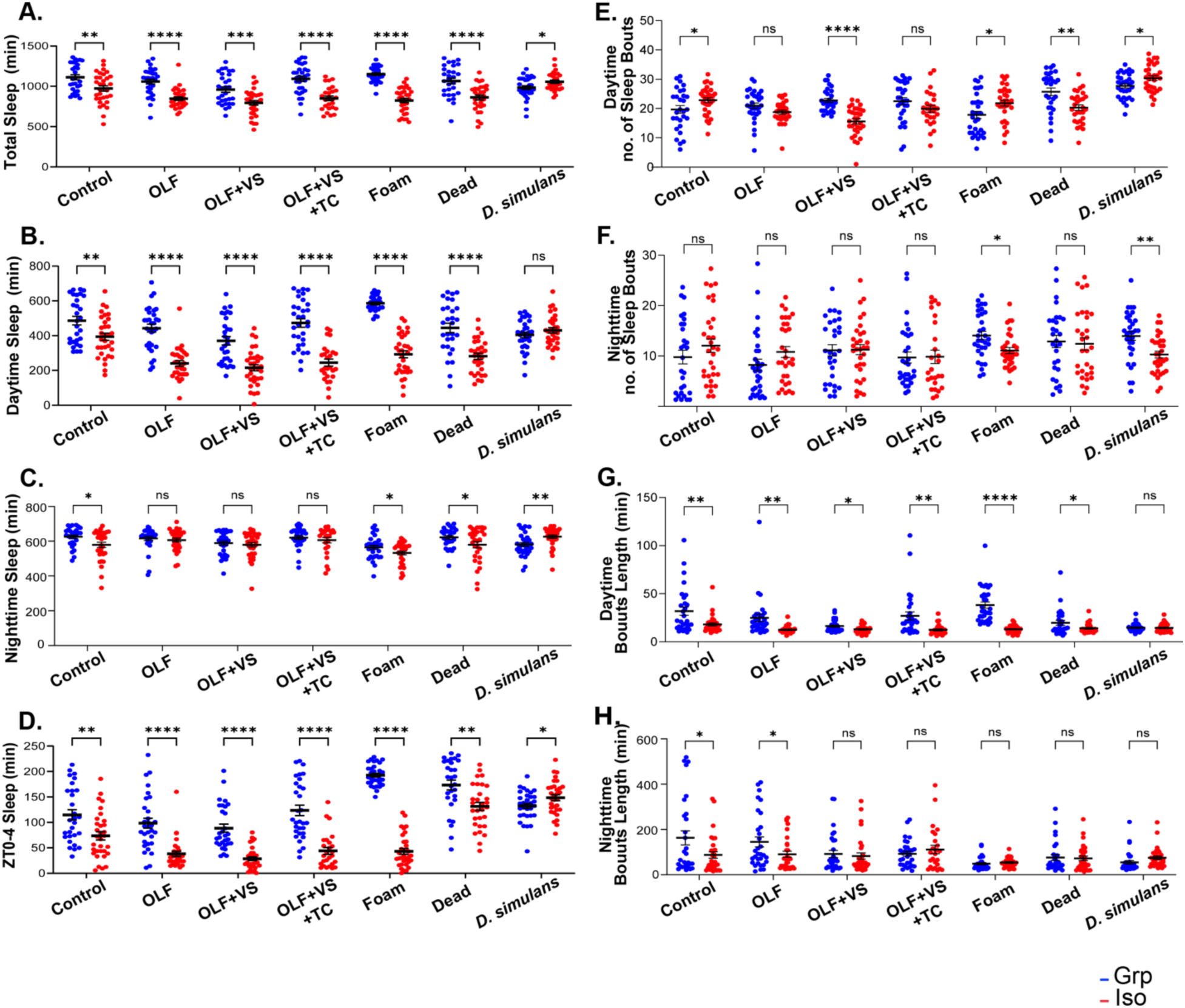
Sensory cues from conspecifics do not alleviate chronic social isolation-induced sleep loss. (A–D) Quantification of total sleep (A), daytime sleep (B), nighttime sleep (C), and ZT0-4 sleep (D). (E–H) Quantification of daytime sleep-bout number (E), nighttime sleep-bout number (F), mean daytime sleep-bout length (G), and mean nighttime sleep-bout length (H). Flies were maintained under the experimental conditions described in Figure 1: Control, OLF (olfactory cues), OLF+VS (olfactory and visual cues), OLF+VS+TC (olfactory, visual, and tactile cues), Foam, Dead conspecifics, and live *D. simulans*. Blue symbols denote grouped flies, and red symbols denote isolated flies. Each dot represents one fly; data are presented as mean ± s.e.m. Statistical analysis was performed using two-sided unpaired t-tests with Welch’s correction. \**P* < 0.05; \*\**P* < 0.01; \*\*\**P* < 0.001; \*\*\*\**P* < 0.0001; NS, not significant; n = 29–32 flies per condition.

**Figure S2.**
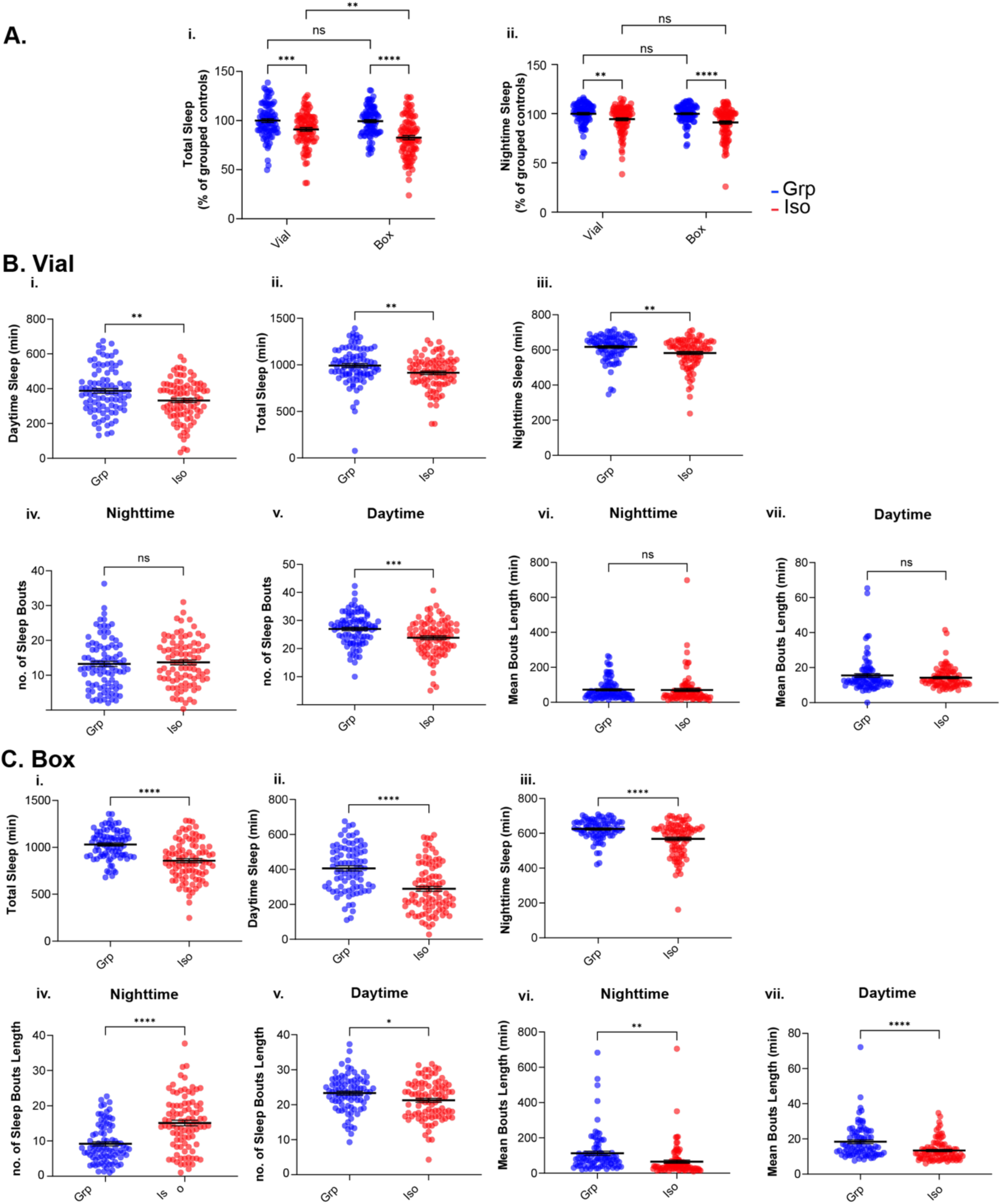
Spatial restriction exacerbates chronic social isolation-induced sleep loss. (A) Quantification of total sleep and nighttime sleep expressed as a percentage of grouped controls, normalized to 100% (mean ± s.e.m.; individual data points shown). Statistical analysis was performed using two-way ANOVA followed by Fisher’s LSD multiple-comparison tests. (B,C) Quantification of all sleep parameters of grouped and isolated animals using (B) vial-based and (C) box-based housing systems. Each dot represents one fly; data are presented as mean ± s.e.m. Statistical analysis was performed using two-sided unpaired t-tests with Welch’s correction. \**P* < 0.05; \*\**P* < 0.01; \*\*\**P* < 0.001; \*\*\*\**P* < 0.0001; NS, not significant; n = 86–92 flies per condition.

**Figure S3.**
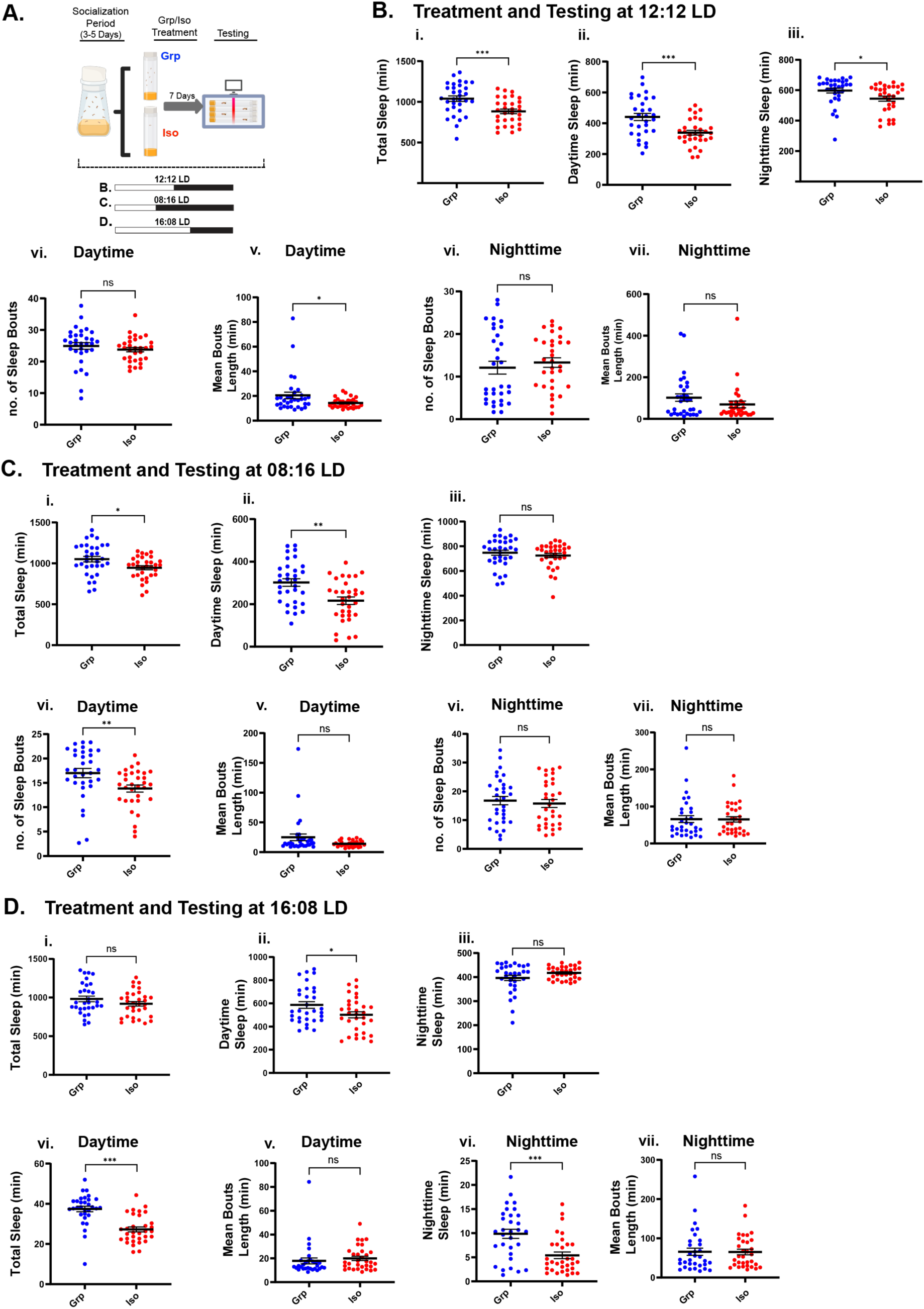
Chronic social isolation alters sleep under different seasonal photoperiods. (A) Schematic of the experimental design. Flies were socialized for 3–5 days and then group-housed (Grp) or isolated (Iso) for 7 days under (B) 12:12 LD, (C) 08:16 LD, and (D) 16:08 LD photoperiods. Following treatment, locomotor activity was recorded under photoperiods matched to those used during treatment. (B–D) Quantification of sleep parameters in grouped and chronically isolated flies under (B) equinox-like 12:12 LD (n = 31–32), (C) winter-like 08:16 LD (n = 32), and (D) summer-like 16:08 LD (n = 31–32) conditions. Data are presented as mean ± s.e.m.; individual data points are shown. Statistical analysis was performed using two-sided unpaired t-tests with Welch’s correction. \**P* < 0.05; \*\**P* < 0.01; \*\*\**P* < 0.001; \*\*\*\**P* < 0.0001; NS, not significant.

**Figure S4.**
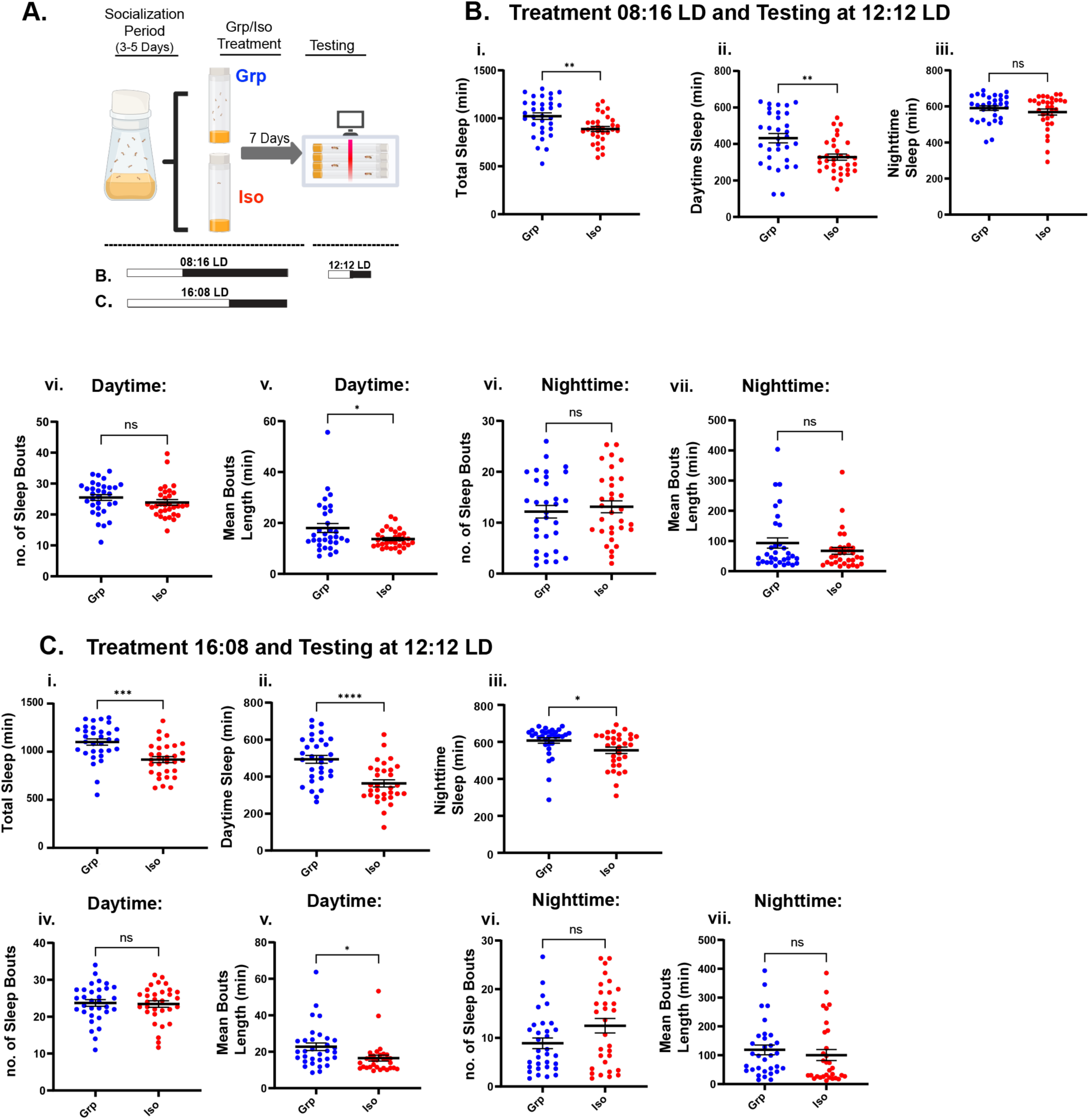
Chronic social isolation alters sleep under different seasonal photoperiods. (A) Schematic of the experimental design. Flies were socialized for 3–5 days and then group-housed (Grp) or isolated (Iso) for 7 days under (B) 08:16 LD or (C) 16:08 LD photoperiods. Following treatment, locomotor activity was recorded under standard 12:12 LD conditions. (B–D) Quantification of total, daytime, and nighttime sleep, as well as daytime and nighttime sleep-bout number and mean bout length, under each treatment photoperiod. Data are presented as mean ± s.e.m.; individual data points are shown. Statistical comparisons were performed using two-sided unpaired t-tests with Welch’s correction. \**P* < 0.05; \*\**P* < 0.01; \*\*\**P* < 0.001; \*\*\*\**P* < 0.0001; NS, not significant; n = 30–32 flies per condition.

**Figure S5.**
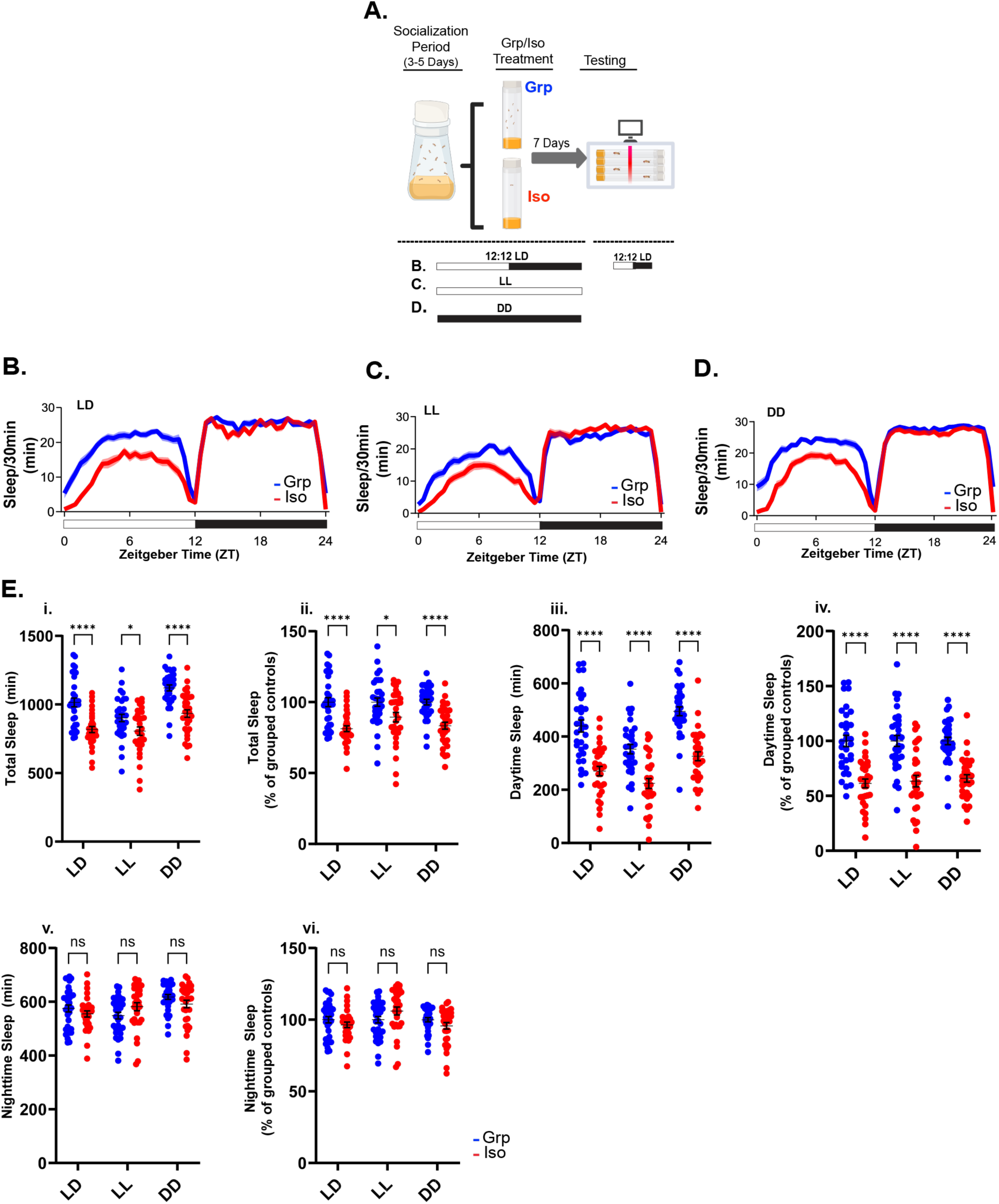
Chronic social isolation reduces sleep independently of environmental light conditions during treatment. (A) Schematic of the experimental design. (B-D) Sleep profiles of group-housed and isolated flies following 3–5 days of socialization and 7 days of group or isolation treatment under (B) 12:12 LD, (C) constant light (LL), or (D) constant darkness (DD), with behavioral testing performed under the 12:12 LD condition. (E) Quantification of daytime, total, and nighttime sleep under each lighting condition, expressed as both a percentage of grouped controls and absolute sleep duration (min): (i–ii) daytime sleep, (iii–iv) total sleep, and (v–vi) nighttime sleep. Data are presented as mean ± s.e.m.; individual data points are shown. Statistical comparisons were performed using two-sided unpaired t-tests. \**P* < 0.05; \*\**P* < 0.01; \*\*\**P* < 0.001; \*\*\*\**P* < 0.0001; NS, not significant; n = 30–32 flies per group.

**Figure S6.**
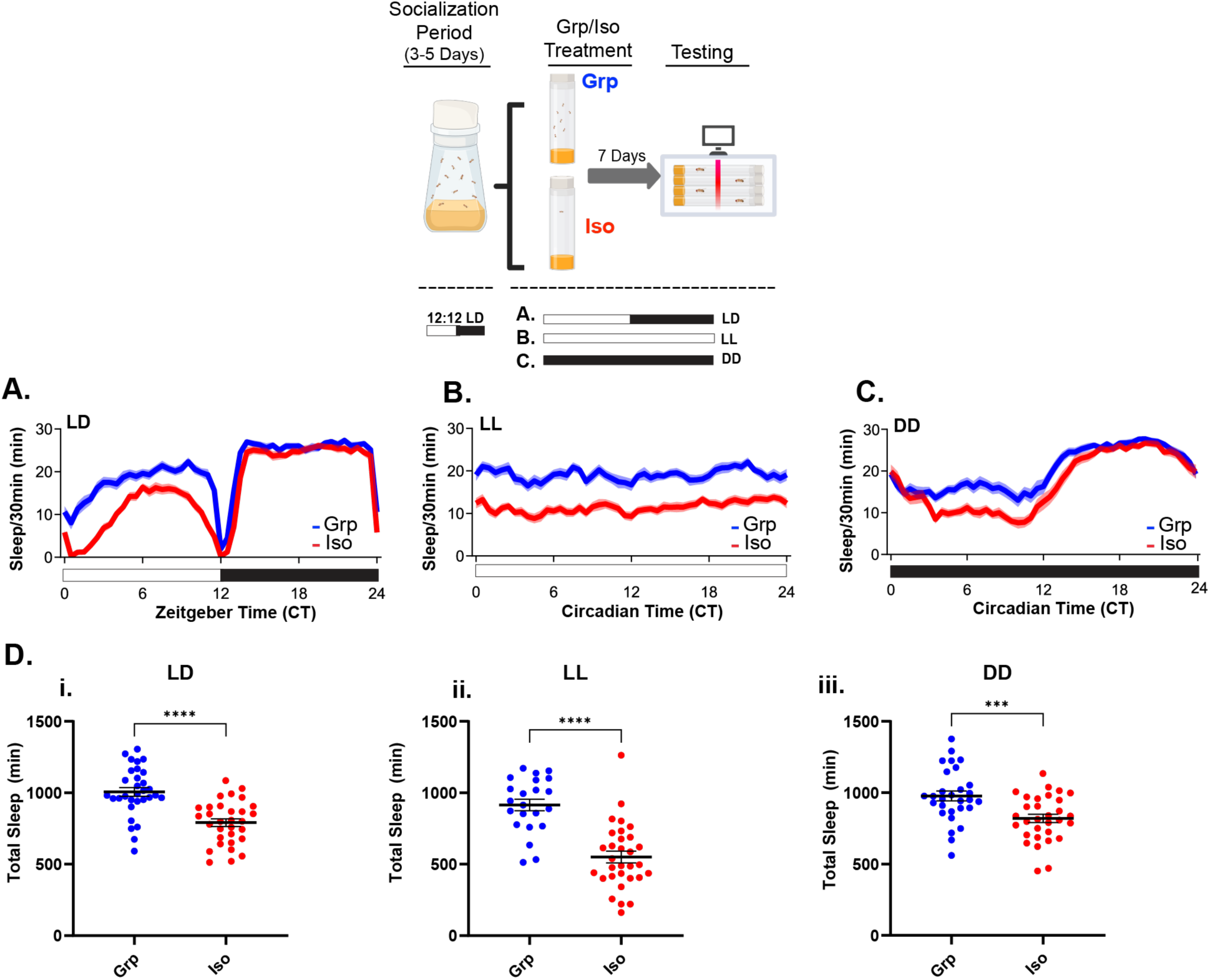
Chronic social isolation reduces sleep independently of environmental light conditions during treatment and testing. (A–C) Schematic of the experimental design. Sleep profiles of group-housed and isolated flies following 3–5 days of socialization and 7 days of group or isolation treatment under (A) 12:12 LD, (B) constant light (LL), or (C) constant darkness (DD), with behavioral testing performed under the corresponding lighting condition. (D) Isolated flies exhibited reduced total sleep under LD (\*\*\*\**P* < 0.0001), LL (\*\*\*\**P* < 0.0001), and DD (\*\*\**P* < 0.001). n = 29–31 flies per condition. Each dot represents one fly; data are presented as mean ± s.e.m. Statistical comparisons were performed using two-sided unpaired t-tests with Welch’s correction.

## References

1. Cacioppo, J.T., et al., Social isolation. Ann N Y Acad Sci, 2011. 1231(1): p. 17–22.

2. Harris, E., Meta-Analysis: Social Isolation, Loneliness Tied to Higher Mortality. JAMA, 2023. 330(3): p. 211–211.

3. Holt-Lunstad, J., Social connection as a critical factor for mental and physical health: evidence, trends, challenges, and future implications. World Psychiatry, 2024. 23(3): p. 312–332.

4. Office of the Surgeon, G., Publications and Reports of the Surgeon General, in Our Epidemic of Loneliness and Isolation: The U.S. Surgeon General’s Advisory on the Healing Effects of Social Connection and Community. 2023, US Department of Health and Human Services: Washington (DC).

5. Steptoe, A., et al., Social isolation, loneliness, and all-cause mortality in older men and women. Proc Natl Acad Sci U S A, 2013. 110(15): p. 5797–801.

6. Stokes, A.C., et al., Loneliness, social isolation, and all-cause mortality in the United States. SSM Ment Health, 2021. 1.

7. Vora, A., et al., The impact of social isolation on health and behavior in Drosophila melanogaster and beyond. Brain Science Advances, 2022. 8(3): p. 183–196.

8. Li, W., et al., Chronic social isolation signals starvation and reduces sleep in Drosophila. Nature, 2021. 597(7875): p. 239–244.

9. LeDoux, J., Rethinking the emotional brain. Neuron, 2012. 73(4): p. 653–76.

10. Pereira, T.D., J.W. Shaevitz, and M. Murthy, Quantifying behavior to understand the brain. Nat Neurosci, 2020. 23(12): p. 1537–1549.

11. Gomez-Marin, A., et al., Big behavioral data: psychology, ethology and the foundations of neuroscience. Nat Neurosci, 2014. 17(11): p. 1455–62.

12. Doria, E.I., et al., GIGEM (Group Isolation Gauge Effect Metrics), a Software Suite for Analyzing Social Isolation-induced Sleep Loss and Multi-batch Experiments in Drosophila. J Biol Rhythms, 2026: p. 7487304261449840.

13. Ganguly-Fitzgerald, I., J. Donlea, and P.J. Shaw, Waking experience affects sleep need in Drosophila. Science, 2006. 313(5794): p. 1775–81.

14. Gil-Marti, B., et al., Socialization causes long-lasting behavioral changes. Sci Rep, 2024. 14(1): p. 22302.

15. Lone, S.R., et al., Social Experience Is Sufficient to Modulate Sleep Need of Drosophila without Increasing Wakefulness. PLoS One, 2016. 11(3): p. e0150596.

16. Donlea, J.M., N. Ramanan, and P.J. Shaw, Use-dependent plasticity in clock neurons regulates sleep need in Drosophila. Science, 2009. 324(5923): p. 105–8.

17. Brown, M.K., E. Strus, and N. Naidoo, Reduced Sleep During Social Isolation Leads to Cellular Stress and Induction of the Unfolded Protein Response. Sleep, 2017. 40(7).

18. MurthyGowda, S., et al., Early life social isolation primes flies for more rapid TDP-43 dependent neurodegeneration later in life. bioRxiv, 2026: p. 2026.09.23.753732.

19. Sun, Y., et al., Social attraction in Drosophila is regulated by the mushroom body and serotonergic system. Nat Commun, 2020. 11(1): p. 5350.

20. Zhao, H., et al., A neural pathway for social modulation of spontaneous locomotor activity (SoMo-SLA) in Drosophila. Proc Natl Acad Sci U S A, 2024. 121(9): p. e2314393121.

21. Sethi, S., et al., Social Context Enhances Hormonal Modulation of Pheromone Detection in Drosophila. Curr Biol, 2019. 29(22): p. 3887–3898.e4.

22. Du, C., et al., Pheromone circuits and transcriptional cascades modulating transcriptional and chromatin states in the Drosophila central brain with social experience. bioRxiv, 2025.

23. Liu, D., et al., A hypothalamic circuit underlying the dynamic control of social homeostasis. Nature, 2025. 640(8060): p. 1000–1010.

24. Hulse, B.K., et al., A connectome of the Drosophila central complex reveals network motifs suitable for flexible navigation and context-dependent action selection. Elife, 2021. 10.

25. Lanz, A.J., et al., Disinhibition of a recurrent attractor gates a persistent goal signal for navigation. bioRxiv, 2025.

26. Dollish, H.K., M. Tsyglakova, and C.A. McClung, Circadian rhythms and mood disorders: Time to see the light. Neuron, 2024. 112(1): p. 25–40.

27. Abrieux, A., et al., EYES ABSENT and TIMELESS integrate photoperiodic and temperature cues to regulate seasonal physiology in Drosophila. Proc Natl Acad Sci U S A, 2020. 117(26): p. 15293–15304.

28. Hidalgo, S., et al., Splicing of a core circadian clock gene regulates seasonal adaptations by a winter gating mechanism. Sci Adv, 2026. 12(29): p. eaed4249.

29. Petsakou, A., T.P. Sapsis, and J. Blau, Circadian Rhythms in Rho1 Activity Regulate Neuronal Plasticity and Network Hierarchy. Cell, 2015. 162(4): p. 823–35.

